# State- and circuit-dependent opponent-processing of fear

**DOI:** 10.1101/2024.05.06.592282

**Authors:** Joanna Oi-Yue Yau, Amy Li, Lauren Abdallah, Leszek Lisowksi, Gavan P. McNally

## Abstract

The presence of valence coding neurons in the basolateral amygdala (BLA) that form distinct projections to other brain regions implies functional opposition between aversion and reward during learning. However, evidence for opponent interactions in fear learning is sparse and may only be apparent under certain conditions. Here we test this possibility by studying the roles of the BLA➔central amygdala (CeA) and BLA➔nucleus accumbens (Acb) pathways in fear learning in male rats. First, we assessed the organisation of these pathways in the rat brain. BLA➔CeA and BLA➔Acb pathways were largely segregated in the BLA but shared overlapping molecular profiles. Then we assessed activity of the BLA➔CeA and BLA➔Acb pathways during two different forms of fear learning - fear learning in a neutral context and fear learning in a reward context. BLA ➔ CeA neurons were robustly recruited by footshock regardless of where fear learning occurred whereas recruitment of BLA➔Acb neurons was state-dependent because footshock only recruited this pathway in a reward context. Finally, we assessed the causal roles of activity in these pathways in fear learning. Photoinhibition of the BLA➔CeA pathway during the footshock US impaired fear learning, regardless of where fear learning occurred. In contrast, photoinhibition of the BLA➔Acb pathway augmented fear learning, but only in the reward context. Taken together, our findings show circuit- and state-dependent opponent processing of fear. Footshock-driven activity in the BLA➔Acb pathway can functionally oppose the BLA➔CeA pathway to limit how much fear is learned.

**Significance Statement:** Here we identify a fear opponent process in the brain. We show that an aversive event can recruit distinct populations of neurons in the rat basolateral amygdala. One population projects to the central amygdala to promote fear learning. A second population projects to the nucleus accumbens to oppose fear learning. These nucleus accumbens projecting neurons limit how much fear is learned and are candidates for therapeutic targeting to minimize the amount of fear learned after a traumatic experience.

---

The basolateral amygdala (BLA) enables learning about emotional events by encoding their valence, assigning that valence to their predictors, and selecting appropriate behavioral responses (e.g. approach versus avoidance) (Janak & Tye, 2015; Tye, 2018). These roles are supported by a complex mosaic of BLA neurons. In mice, BLA principal glutamatergic neurons can be distinguished on the basis of their long-range projections. BLA neurons projecting to the central amygdala (CeA) respond preferentially to aversive events (e.g. quinine, footshock) and their predictors whereas BLA neurons projecting to the nucleus accumbens (Acb) respond preferentially to rewarding events (e.g. sucrose) and their predictors (Beyeler et al., 2016). BLA principal neuron molecular heterogeneity overlaps with these circuit features and may contribute to valence and learning. For example, BLA principal neurons expressing *Ppp1r1b* have been linked to positive valence whereas neurons expressing *Rspo2* have been linked to negative valence (Kim, Pignatelli, Xu, Itohara, & Tonegawa, 2016). There are important nuances within this general BLA mosaic. For example, a subpopulation of *Rspo2* BLA neurons expressing *Fezf2* encode positive versus negative valence based on their long-range projections to the nucleus accumbens and olfactory tubercle (Zhang et al., 2021).

The presence and organisation of BLA valence coding circuits is suggestive of opponent processing of learning, where an opponent (e.g., reward or relief) process is recruited during learning to limit how much fear is learned (Konorski, 1967; Schull, 1979; Solomon & Corbit, 1974). This possibility is supported by findings that positive and negative valence BLA neurons can mutually inhibit each other, most likely via parvalbumin interneurons (Kim et al., 2016; Piantadosi et al., 2024). However, despite compelling evidence for opponent processes at the behavioral level, as shown via phenomena such as superconditioning and counterconditioning (Dickinson & Dearing, 1979; Nasser & McNally, 2012, 2013), evidence for opposition between BLA circuits during fear learning is sparse. We do know that the BLA➔CeA pathway is important for fear learning because optogenetic inhibition of BLA➔CeA neurons impairs fear learning (Namburi et al., 2015). However, optogenetic inhibition of the reward coding BLA➔Acb pathway, which an opponent process model predicts should augment fear learning, had no effect on fear learning (Namburi et al., 2015).

So, if there is functional opposition between BLA aversion and reward coding circuits, this opposition may only be apparent under certain conditions. Here, we test these ideas by studying the roles of the BLA ➔ CeA and BLA ➔ Acb pathways in fear learning in rats. First, we assessed the anatomical and molecular organisation of these pathways in the rat brain. Then we assessed activity of these pathways during two different forms of fear learning - fear learning in a neutral context and fear learning in a reward context. Finally, we assessed the causal roles of activity in these pathways in fear learning in a neutral context and a reward context. Our findings show, for the first time, circuit- and state-dependent opponent processing of fear. Footshock-driven activity in the BLA ➔ Acb pathway can functionally oppose the BLA ➔ CeA pathway to limit how much fear is learned.

## Methods

### Subjects

Subjects were experimentally naive male Long Evans rats (280 – 500g) obtained from the Animal Resource Centre (Perth, Australia) and from The University of New South Wales. Female rats were not available and we consider this limitation in the Discussion. Rats were housed in groups of 2 – 4 in a climate-controlled colony room. Lights were on at 7:00AM and off at 7:00PM, with all experiments conducted in the light cycle. Animals were allowed free access to chow and water unless otherwise stated.

### Behavioural apparatus

Behavioural training was conducted in identical Med-Associate chambers. The chambers were 24cm (length) x 30 cm (width) x 21 cm (height) and enclosed in a ventilated, sound-attenuating cabinets measuring 59.5 cm (length) x 59 cm (width) x 48 cm (height). The left sidewall was fitted with a magazine dish where grain pellets (Bio-Serv, USA) were sometimes delivered when a retractable lever located 4 cm to the right of the magazine was pressed. A 3 W house light was mounted on top of the right sidewall next to a speaker used to delivery auditory CSs. A metal grid was fitted to the floor of the chamber to deliver a scrambled footshock US. For optogenetics experiments, an LED with an integrated rotary joint (Doric Instruments, Canada) was suspended above the chamber and controlled by an LED driver connected to MED-Associates.

### Retrograde tracers and viral vectors

Cholera toxin subunit B (recombinant) (CTb) Alexa Fluor 488 (Cat#C34775, Therma Fisher Scientific, USA) and CTb Alexa Fluor 555 (Cat#CC34776, Therma Fisher Scientific, USA) were used for anatomical and molecular investigation of BLA output pathways. AAVs driven from a CamKIIα promoter were used to globally target BLA neurons whereas retrograde AAVs were used to target specific BLA pathways. We used AAV5-CaMKIIα-eNpHR3.0-eYFP (6×10^12^) and AAV5-CaMKIIα-eYFP (4×10^12^) (both from UNC Vector Core) to target BLA neurons. We packaged pAAV-hSyn-eNpHR 3.0-EYFP (a gift from Karl Deisseroth [Addgene plasmid #26972; http://n2t.net/addgene:26972; RRID:Addgene_26972]) and pAAV-hSyn-EGFP (a gift from Bryan Roth [Addgene plasmid # 50465; http://n2t.net/addgene:50465;RRID:Addgene_50465]) into a retrograde AAV helper vector (a gift from Alla Karpova & David Schaffer [Addgene plasmid # 81070; http://n2t.net/addgene:81070; RRID:Addgene_81070]) (Tervo et al., 2016), at the Vector Genome Facility (Westmead Children’s Hospital, Sydney, Australia) to obtain AAV2_retro_-hSyn-eNpHR3.0-eYFP (2.96×10^13^) and AAV_retro_-hSyn-eGFP (1×10^13^). We used AAVretrograde-gCaMP7f (1.8×10^13^) (pGP-AAV-syn-jGCaMP7f-WPRE was a gift from Douglas Kim & GENIE Project [Addgene viral prep # 104488-AAVrg; http://n2t.net/addgene:104488; RRID:Addgene_104488) and AAVretrograde-jRGECO1a (7×10^12^) (pAAV.Syn.NES-jRGECO1a.WPRE.SV40 was a gift from Douglas Kim & GENIE Project (Addgene viral prep # 100854-AAVrg; http://n2t.net/addgene:100854; RRID:Addgene_100854) (Dana et al., 2016).

### Surgery

Rats were anesthetised with isoflurane (5% induction; 2% maintenance) mixed with oxygen and placed in a stereotaxic frame (Kopf Instruments, CA, USA). Prior to incision, rats received the analgesic carprofen (Rimadyl, Zoetis; 5mg/kg) via a subcutaneous injection and 0.5% bupivacaine under the incision site. Following an incision to expose the skull, a hand drill was used to make a craniotomy above each injection and cannula implantation site. A 23-gauge, cone-tipped 5ml stainless-steel injector (SGE Analytical Sciences) holding tracer or AAV was lowered into each injection site and infused at a rate of 100nl/min (UltraMicro Pump III with SYS-Micro4 Controller, World Precision Instructions, USA). For CeA these co-ordinates were −2.25 (AP), ±4.25 (ML) and −7.9 (DV). For Acb, these co-ordinates were +1.2 to 1.3 (AP), ±1.5 or ±1.7 at 6° (ML) and 7.5 (DV). For BLA, these coordinates were −3.24 (AP), ±5.15 (ML), and −8.1 (DV). All coordinates are in mm from Bregma. The injector was left in place for additional 7 mins for diffusion. For photometry and optogenetics experiments, a 400μm fibre optic cannula was implanted unilaterally (photometry) or bilaterally (optogenetics) above the BLA (AP: −3.24; ML: ±5.1, DV: −7.9, in mm from Bregma) and secured in place by dental cement (Vertex Dental) anchored to jeweller’s screws attached to the skull. The incision was sutured and antibiotic (Duplocillin, Intervet; 40,000-60,000 IU/kg) was given intraperitoneally. Rats were monitored until the end of the experiment.

We used CTb Alexa Fluor 488 or Alexa Fluor 555 conjugate to label CeA- and Acb-projecting BLA cells in the same rat. One variant was injected into the CeA and the other was injected into the Acb of the same hemisphere. Each injection was 200nl in volume. The tracer colour variant and the injected hemisphere were counterbalanced across rats.

We used unilateral injection of the AAV containing CaMKIIα-gCaMP7f (fibre photometry) or bilateral injection of the AAV-CaMKIIα-eNpHR3.0-eYFP or the control AAV-CaMKIIα-eYFP (optogenetics) to target BLA glutamatergic neurons non-selectively. Each injection was 750nl.

We used retrograde AAVs containing the green (gCaMP7f) (Dana et al., 2019) or red-shifted (jRGECO1a) (Dana et al., 2016) calcium sensor to target BLA output pathways for simultaneous recording. Each rat received three craniotomies in the same hemisphere – one above the CeA, one above the Acb and one above the BLA. One sensor was injected into the CeA at a volume of 300nl, and the other was injected into the AcbSh at a volume of 500nl. The sensor used at each projection site and the hemisphere injected was counterbalanced.

We used retrograde AAVs containing either eNpHR3.0 or the control eGFP to photoinhibit the BLA➔CeA and BLA ➔ Acb pathways. Rats received bilateral AAV injections of into the CeA (100nl injection) or Acb (750nl injection).

### Single-molecule FISH

Brains were extracted 7 days after CTb injections, snap-frozen with liquid nitrogen, and stored at −80°C. Brains were sliced coronally at 10 µm and the BLA was collected onto superfrosted microscope slides. Accumbens and amygdala regions were also collected on separate slides to confirm CTb injection sites. We used an RNAScope Fluorescent HiPlex Kit (Advanced Cell Diagnostics) with *Rspo2* (#ADV1190711T6), *Ppp1r1b* (#ADV1048941T9) and *Fezf2* (#ADV1190731T12) probes custom designed for rat. BLA samples were treated with paraformaldehyde, dehydrated with ethanol and treated with Protease IV for 20mins. Slides were incubated with target probes and each signal was sequentially amplified using HiPlex Amp 1-3. Dylight650 was applied to first label *Rspo2* and counterstained with DAPI. Slides were coverslipped and DAPI, CTb 488, CTb555 and Dylight650 channels were imaged using an Axioscan 7 slide scanner (Zeiss, Germany). The next day, coverslips were removed and Dylight650 was cleaved and reattached to label *Ppp1r1b* and the 4 channels were reimaged. This process was repeated once more to label and image *Fezf2*-expressing cells. Images were then collated using ZEN software (Zeiss, Germany) and labelled cells in the BLA were counted using CellProfiler (Stirling et al., 2021).

### Behavioural procedures

#### Fibre photometry

Rats were food restricted to ∼ 85% of their body weight 2 days prior to behavioral training and they were maintained on this feeding regime until the end of the experiment. Rats received training in two separate contexts each day. In one context (reward context), a lever was available to earn a food pellet reward delivered into a magazine. In the second context (neutral context), a slate was slotted into the wall to block access to the magazine and to the lever slot. These contexts were further distinguished by olfactory (peppermint, rosewater), auditory (fan on or off), visual (houselight on or off) as well as spatial (different box with same dimensions in a different location) cues which were counterbalanced across rats.

On Day 1 – 5, rats received daily 1 hr lever press training in the reward context and were placed in neutral context for the same duration. Context order was randomised each day. On Day 1 – 2, rats received magazine training in the reward context. In these sessions, every lever press was rewarded by grain pellet delivery, with additional ‘free’ grain pellets delivered on FI300 schedule. Sessions were terminated when the rat reached 100 lever presses or after 60 min had elapsed. On Day 3, rats lever pressed for pellets on a VI30 schedule and on Days 4 and 5 on a VI120 schedule. On Day 6, rats were pre-exposed to one auditory CS in the neutral context, and a different auditory CS in the appetitive context where rats were lever pressing on a VI120 schedule. Auditory CSs were either a 60s 10Hz clicker or 60s tone counterbalanced across context and rats. In each context, the 60s CS was presented 4 times on an ITI of 600 – 900 s during a 1 hr session. On Day 7, rats were tethered to patch cables and then subjected to fear conditioning in both contexts. In each context, the CS was presented 4 times, and each presentation co-terminated with the 0.5mA, 0.5s footshock US during the 1 hr session.

Recordings were made using the Fibre Photometry System from Doric Lenses and Tucker Davis Technologies (RZ5P, Synapse). For CaMkIIα-gCaMP recordings, 465nm and 405 nm wavelength light was emitted from LEDs controlled by programmable LED drivers. For dual-colour, simultaneous recordings of BLA output pathways, an additional 560 nm wavelength light was emitted. Excitation lights were channelled through patch cables (0.39NA, Ø400mm core multimode prebleached patch cables) and light intensity at the tip of the cable was maintained at 10-13 µW across sessions. Ca^2+^-dependent and isosbestic fluorescence were amplified and measured by Doric Fluorescence Detectors. Synapse software controlled and modulated excitation lights (465 nm, 209Hz; 405 nm, 331 Hz; 560nm, 537Hz) as well as demodulated and low-pass filtered (3 Hz) transduced fluorescence signals in real time via the RZ5P. RZ5P/Synapse also received Med-PC signals to record behavioural events in real time.

#### Photoinhibition during fear learning

To assess the roles of BLA and BLA output pathways in fear learning in the reward context, rats were first restricted to ∼85% of their body weight 2 days prior to behavioural training and were maintained on this schedule until the end of behavioural experimentation. Rats received one training session per day. On Day 1 – 2, rats received magazine training as described above. On Day 3, rats lever pressed for pellets on a VI30 schedule in a 2-hr session. From Day 4 onwards, rats were maintained on a VI120 schedule during daily 2 hr sessions. From Day 7 onwards, rats were tethered to patch cables. Rats received pre-exposure to the auditory CS (85dB 10Hz clicker) on Days 9 – 10 whilst lever pressing. In each 2-hr session, rats received 4 60s presentations of the clicker CS with an ITI between 1200 – 1800 s. Rats were fear conditioned to the CS on Day 11 – 13. Prior to each 2-hr session, rats were tethered to patch cables connected to a 625nm LED controlled by an LED driver programmed via Med-Associates. Rats received 4 presentations of the CS on a randomised ITI ranging between 1200 – 1800 s. Each CS co-terminated with a 0.5mA, 0.5s footshock US. Photoinhibition occurred at the time of US onset and continued for 4.5s after US offset. Rats were tested for their fear on Day 14 – 17. In each 70 min test session, rats were presented with 4 non-reinforced presentations of the CS on an ITI of 900 s.

To assess the role of the BLA➔Acb pathway in fear learning in the neutral context, on Day 1, animals were pre-exposed to the shock context for 10 min while tethered to dummy patch cables. No stimuli were presented. On Day 2, rats underwent a 20 min fear conditioning involving three presentations of the 60 second auditory CS (85dB 10Hz clicker) at 120s ITI that either co-terminated with a 0.5 s 0.35 mA or 0.7 mA footshock US. Photoinhibition occurred at the time of US onset and continued for 4.5s after US offset US onset and for 4.5 seconds thereafter, animals received constant optical stimulation of 625 nm wavelength light. On Days 3 and 4, rats were tethered to dummy patch cables and tested for their fear to the CS in 20 min sessions. They received six non-reinforced presentations of the 60 second CS at a ITI of 120 seconds.

### Histology

For retrograde tracing studies, rats were perfused 10 days after surgery, brains were extracted and sliced coronally at 40 µm using a cryostat (Lecia Microsystems). The BLA was collected onto slides, dried and coverslipped. Four BLA sections were examined from each animal: one anterior (−2.40 mm to −2.64 mm from bregma), one middle (−2.92 mm to −3.12 mm from bregma), and two posterior (−3.36 mm to −3.48 mm, and at −3.60 mm from bregma). The CeA and Acb were also collected and assessed to verify injection sites. Two-channel fluorescent images of the BLA were obtained using an Axioscan 7 slide scanner (Zeiss, Germany). For single molecule FISH, three sections were analysed from each animal: one middle (−2.92 mm to −3.12 mm from bregma), and two posterior (−3.36 mm to −3.48 mm, and at −3.60 mm from bregma).

gCaMP7f and jRGECO1a expression were verified using two-colour fluorescence. Free-floating tissue was washed in phosphate buffer (PB) and non-specific binding was blocked using a mixture of 2.5% normal donkey serum (NDS) and normal goat serum (NGS) in PB. Tissue was incubated in 1:1500 chicken anti-GFP (Thermo Fisher Scientific Cat# A10262, RRID:AB_2534023) and rabbit anti-RFP (Thermo Fisher Scientific Cat# 710530, RRID:AB_2532732) diluted in PBTX and 2% NGS and 2% NDS overnight in room temperature. Unbounded primary antibodies were washed off and then incubated in 1:1000 Alexa 488 goat anti-chicken (Thermo Fisher Scientific Cat# A-11039, RRID:AB_2534096) and 1:1000 Alexa 594 donkey anti-rabbit (Thermo Fisher Scientific Cat# A-21207, RRID:AB_141637) for 4h in room temperature. Sections were washed and mounted onto frosted microscope slides and coverslipped with Permafluor (Thermo Fisher Scientific, USA). eGFP and eNpHR3.0 expression were revealed using diaminobenzidine (DAB) immunochemistry. Free-floating tissue was washed and dehydrated in alcohol and non-specific binding was blocked (5% NHS in PB). Sections were incubated in 1:2000 rabbit anti-GFP (Thermo Fisher) diluted in PBTX and 2% NHS for 24h in room temperature. Sections were washed and incubated in 1:2000 biotinylated donkey anti-rabbit (Jackson ImmunoResearch Laboratories) diluted in PBTX and 2% NHS overnight in room temperature. Sections were washed and incubated in avidin-biotin complex diluted in PBTX for 2hr in room temperature. Sections were washed and incubated in DAB solution for 15 minutes. Reactions were initiated by 0.2µl/ml glucose oxidase aspergillus (Sigma-Aldrich) and stopped by washing sections with acetate buffer. Sections were washed and mounted onto gelatin-coated microscope slides. Slides were dehydrated in ethanol and cleared in histolene and then coverslipped with Entellan (ProSciTech, Australia). For retrograde tracing verification, placements were confirmed using native fluorescence viewed under an Olympus BX52 fluorescent microscope.

### Statistics and data analyses

Fear was measured via freezing in the neutral context and via conditioned suppression of lever pressing in the reward context. Freezing (the absence of all movement except that required for breathing) (Fanselow, 1994) was scored every 2 seconds and the number of observations scored as freezing was converted to a percentage. Suppression ratios were calculated as CS lever presses/(PreCS lever presses + CS lever presses), with pre-CS defined as the 1 min prior to CS onset. A ratio of 0.5 indicates no suppression, as there are the same number of lever presses during the CS as well as the minute prior. A ratio of 0 indicates complete suppression of lever pressing during the CS. These data were analysed via repeated measures ANOVA.

Fibre photometry signals were extracted to MATLAB and down sampled (15.89 Hz). Each signal (isosbestic 405 nm, Ca^2+^-dependent 465 nm, and Ca^2+^-dependent 560nm during dual colour photometry) was high-pass (90 s) and low-passed (1 Hz) filtered. Robust regression (Keevers, McNally, & Jean-Richard-Dit-Bressel, 2024) was used to fit the isosbestic signal onto Ca^2+^-dependent signals, and a separate DF/F was calculated for gCaMP (green) and jRGECO1a (red) via (Ca2+ dependent signal – fitted 405nm)/fitted 405nm. DF/F signals around CS and US onset were isolated and aggregated; the 3 s before each event was used as a baseline, and the 7 s following each event was defined as the event transient. A bootstrapping confidence interval (CI) procedure (95% CI, 1000 bootstraps) was used to determine significant event-related transients within this window (Jean-Richard-dit-Bressel, Clifford, & McNally, 2020). A distribution of bootstrapped DF/F means was generated by randomly resampling from trial DF/F waveforms, with replacement, for the same number of trials. A confidence interval was obtained per time point using the 2.5 and 97.5 percentiles of the bootstrap distribution, which was then expanded by a factor of sqrt(n/(n-1)) to adjust for narrowness bias. Significant transients were defined as periods where 95% CI did not contain zero (baseline) for at least 0.5 s. Cross-correlation analyses were conducted using MATLAB with dF/F signals around US onset (−3 to 7 s).

## Results

### Basolateral amygdala pathways are segregated

First, we investigated the organisation of BLA pathways to the Acb and CeA. We injected two differently coloured retrograde tracers (N = 6), Alexa Fluor 488 and Alexa Fluor 555, into Acb and CeA and mapped retrograde labelled neurons in the BLA (Figure 1A and B). The vast majority (>97%) of labelled cells were single labelled, expressing only one of the two tracers, and there were significantly more single labelled compared to double labelled (∼3%) cells (F(1,5) = 97.05, p < .001; Figure 1C and D), confirming that these two pathways are mostly segregated in the rat, consistent with previous findings from mice (Beyeler et al., 2018). CeA- and Acb-projecting neurons were intermingled and distributed across the anterior-posterior axis of the BLA (Figure 1D). They were also distributed across BLA subnuclei, but distinct subpopulations were preferentially located in different nuclei (Figure 1E – G). CeA-projecting neurons were distributed across the rat BLA, with no differences across the lateral amygdala (LA), basolateral amygdala (BL) or basomedial amygdala (BM), largest F(1,5) = 4.30, p > .05). Meanwhile, there were more Acb-projecting neurons in the BL (F(1,5) = 37.79, p = .002) and BM (F(1,5) = 24.44, p = .004) than in the LA, and more Acb-projecting neurons in the BL than the BM (F(1,5) = 9.62, p = .027). For dual-labelled neurons, there was a greater percentage of neurons in the BL relative to the LA (F(1,5) = 20.89, p = .006).

**Figure 1.**
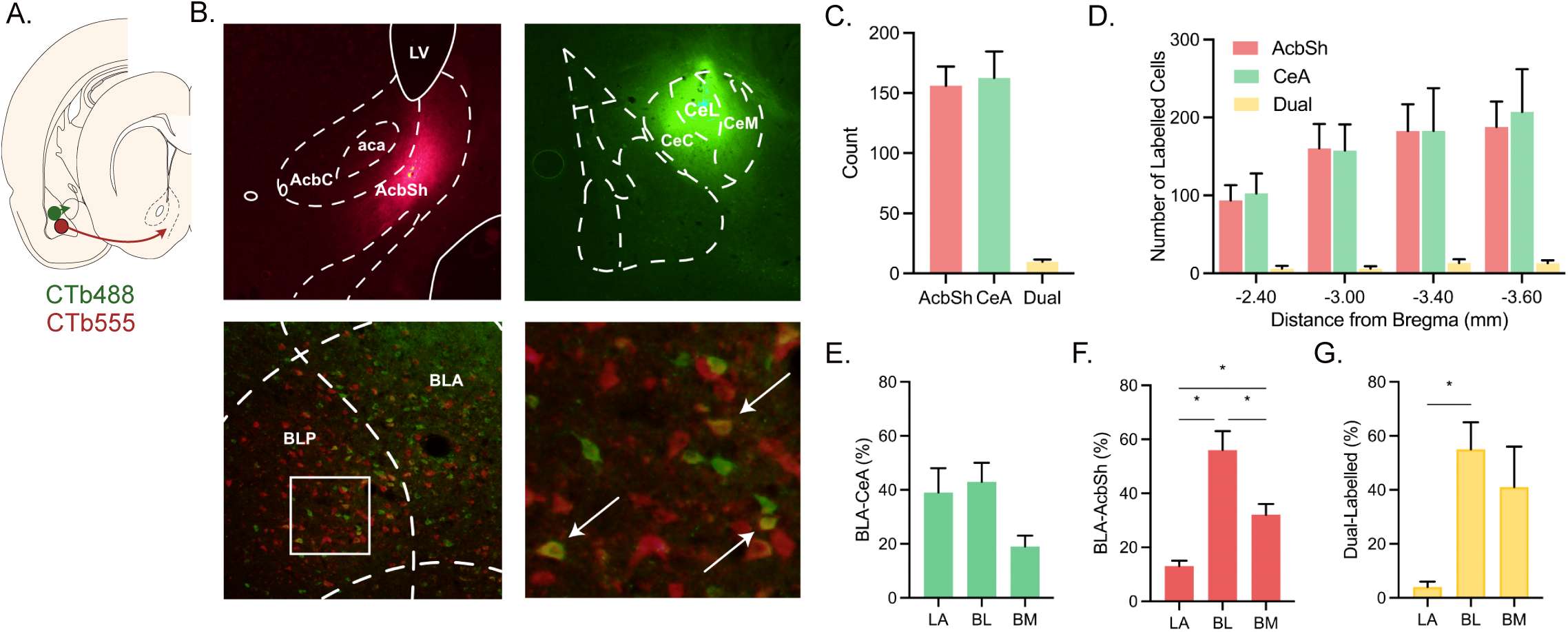
Segregation of basolateral amygdala output pathways. A) CTb-488 and CTb-555 were applied to central amygdala (CeA) or nucleus accumbens (Acb) and retrograde labelled neurons in the basolateral amygdala (BLA) were identified. B) Representative CTb deposits in the Acb, CeA and retrograde labelled neurons in the BLA. C) Counts of single and dual labelled neurons in the BLA. D) Anterior-posterior distribution of single and dual labelled neurons in the BLA. E) – G) Amygdala subnucleus distribution of single and dual labelled neurons.

### Basolateral amygdala pathways share molecular profiles

Different functional BLA neuron populations have been distinguished based on expression of distinct genetic markers in mice, most notably the genes *Rspo2*, *Ppp1r1b* and *Fezf2* (Kim et al., 2016; Zhang et al., 2021). So, we asked whether these markers also distinguish between the BLA ➔ CeA and BLA ➔ Acb pathways in rats. We injected retrograde tracers into the rat (N = 4) CeA and Acb and used single-molecule FISH to label BLA for *Rspo2*, *Ppp1r1b* and *Fezf2* mRNA expression (Figure 2A – C). Approximately half of all BLA cells expressed at least one of these mRNA targets (Figure 2D), confirming their abundant expression in the rat BLA. Interestingly, *Rspo2*, *Ppp1r1b* and *Fezf2* were expressed at similar levels (F(1,3) = 6.725, p > 0.05; Figure 2E). Of note, most labelled cells were tripled labelled with all 3 gene targets (Figure 2F). We then asked whether these gene expression profiles related to BLA ➔ CeA and BLA ➔ Acb pathways (Figure 2G). We found the expression profiles of *Rspo2*, *Ppp1r1b* and *Fezf2* were similar across the BLA ➔ CeA and BLA ➔ Acb pathways (Figure 2H and I). A majority of neurons at the origin of each pathway expressed all three mRNAs (He, Huang, Schlüter, & Dong, 2023; Kim et al., 2017; Madur et al., 2023). So, although *Rspo2*, *Ppp1r1b* and *Fezf2* expression can distinguish valence coding neurons in the mouse BLA (Kim et al., 2016; Zhang et al., 2021), these three genes are highly co-localised in rat BLA and do not distinguish between the BLA➔CeA and BLA➔Acb pathways.

**Figure 2.**
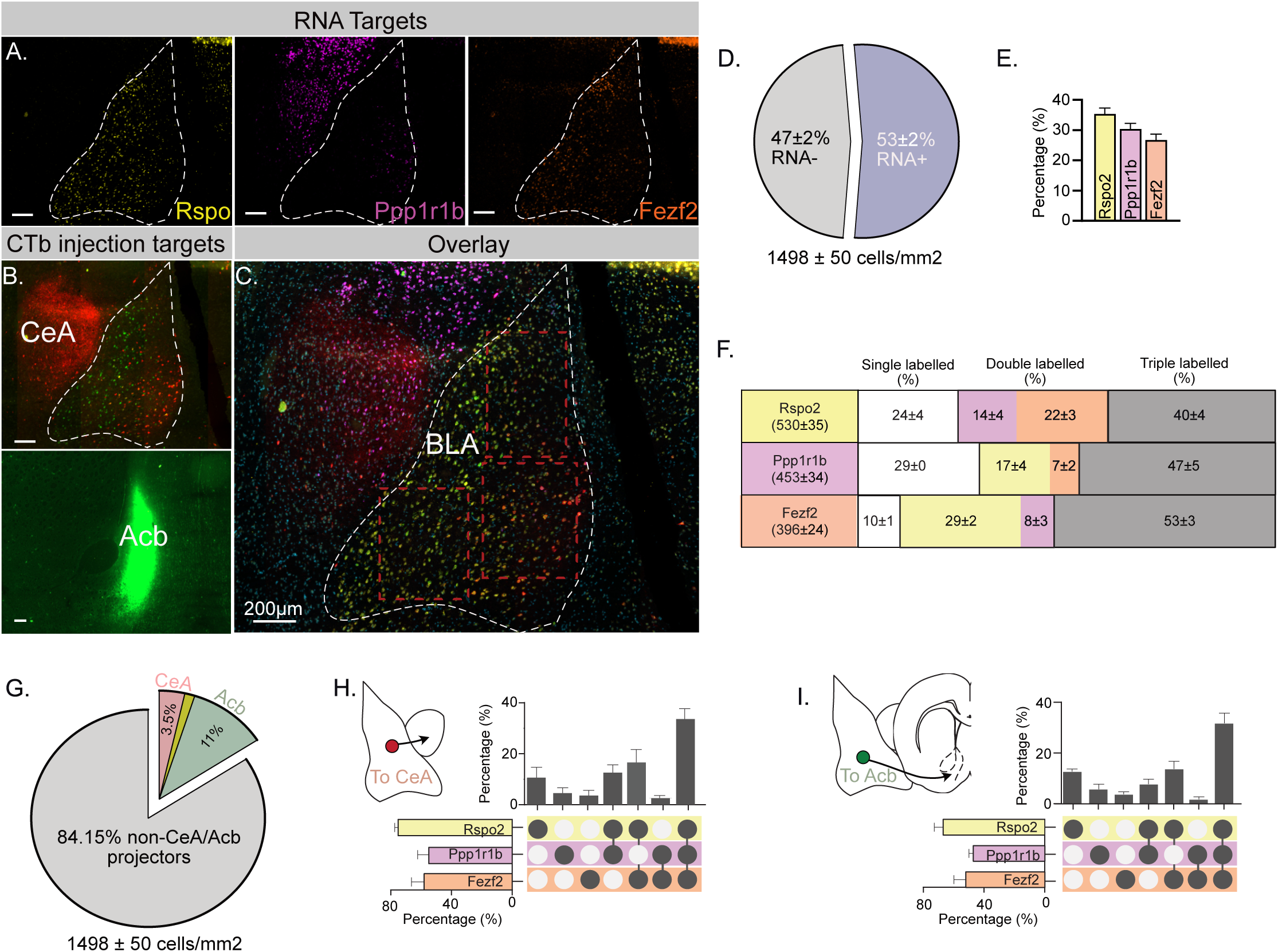
Molecular characterization of basolateral amygdala output pathways. A) Representative photomicrographs showing distribution of *Rspo*, *Ppp1r1b*, and *Fezf2* mRNA in rat BLA. B) Representative CTb deposits in the Acb, CeA; C) Overlay of retrograde labelled neurons and *Rspo*, *Ppp1r1b*, and *Fezf2* mRNA in rat BLA. D) Percentage of BLA neurons expressing any of *Rspo*, *Ppp1r1b*, and *Fezf2* mRNA. E) Percentage of BLA neurons expressing either *Rspo*, *Ppp1r1b*, or *Fezf2* mRNA. F) Percentage of BLA neurons single, double, or triple labelled for *Rspo*, *Ppp1r1b*, or *Fezf2* mRNA. G) Percentages of BLA ➔ CeA and BLA ➔ Acb neurons. single, double, or triple labelled for *Rspo*, *Ppp1r1b*, or *Fezf2* mRNA. Scale bars = 200μm.

### BLA output pathways show circuit- and state-dependent US responsivity

The BLA has important roles in both positive and negative valence. These roles can be distinguished by different BLA circuits, with BLA➔CeA pathway important for negative valence and BLA➔Acb pathway important for positive valence (Beyeler et al., 2018, 2016; Namburi et al., 2015). This raises the possibility that these two pathways may act in opposition to each other during fear learning. However, whether these two BLA circuits have the same or different activity profiles during fear learning is unknown. Furthermore, whether the conditions under which fear is learned determines these activity profiles is unknown.

To answer these questions, we first injected CaMKIIα-gCaMP7f into the rat BLA (N = 8) and used fibre photometry to record from a global population of BLA neurons (Figure 3A) while the same rats underwent two different forms of fear conditioning. In the neutral context, the conditioning chamber was bare except for the wall walls, roof, and grid floor and rats received pairings of an auditory conditioned stimulus with a 0.5mA, 0.5s footshock unconditioned stimulus (US). Conditioned freezing was measured as the fear response to presentations of the auditory conditioned stimulus. In the reward context, a lever was available and lever pressing led to a scheduled delivery of a food pellet to a magazine. Pairings of the auditory conditioned stimulus with a 0.5mA, 0.5s footshock US were superimposed on this lever pressing in a response-independent manner. Conditioned suppression of lever pressing was measured as the fear response because rats express little to no freezing under these conditions. It is important to note that the contexts were matched on the total time rats had spent in them prior to and during conditioning. Rats acquired fear to both auditory CSs (Figure 3B and C). In the neutral context, rats increased freezing to the auditory CS across training (main effect of Trial: F(1,7) = 12.222, p < 0.01). Likewise in the reward context, lever pressing rates (as shown by suppression ratios) decreased during the different auditory CS across training (main effect of Trial: F(1,7) = 53.226, p < 0.01). BLA neurons showed excitatory Ca^2+^- transients to the auditory CS and the footshock US in both neutral and reward contexts (Figure 3D and E). In this and remaining figures, periods of statistical significance in photometry data are shown by coloured bars above the waveform indicating times when 95% confidence interval does not include 0% DF/F.

**Figure 3.**
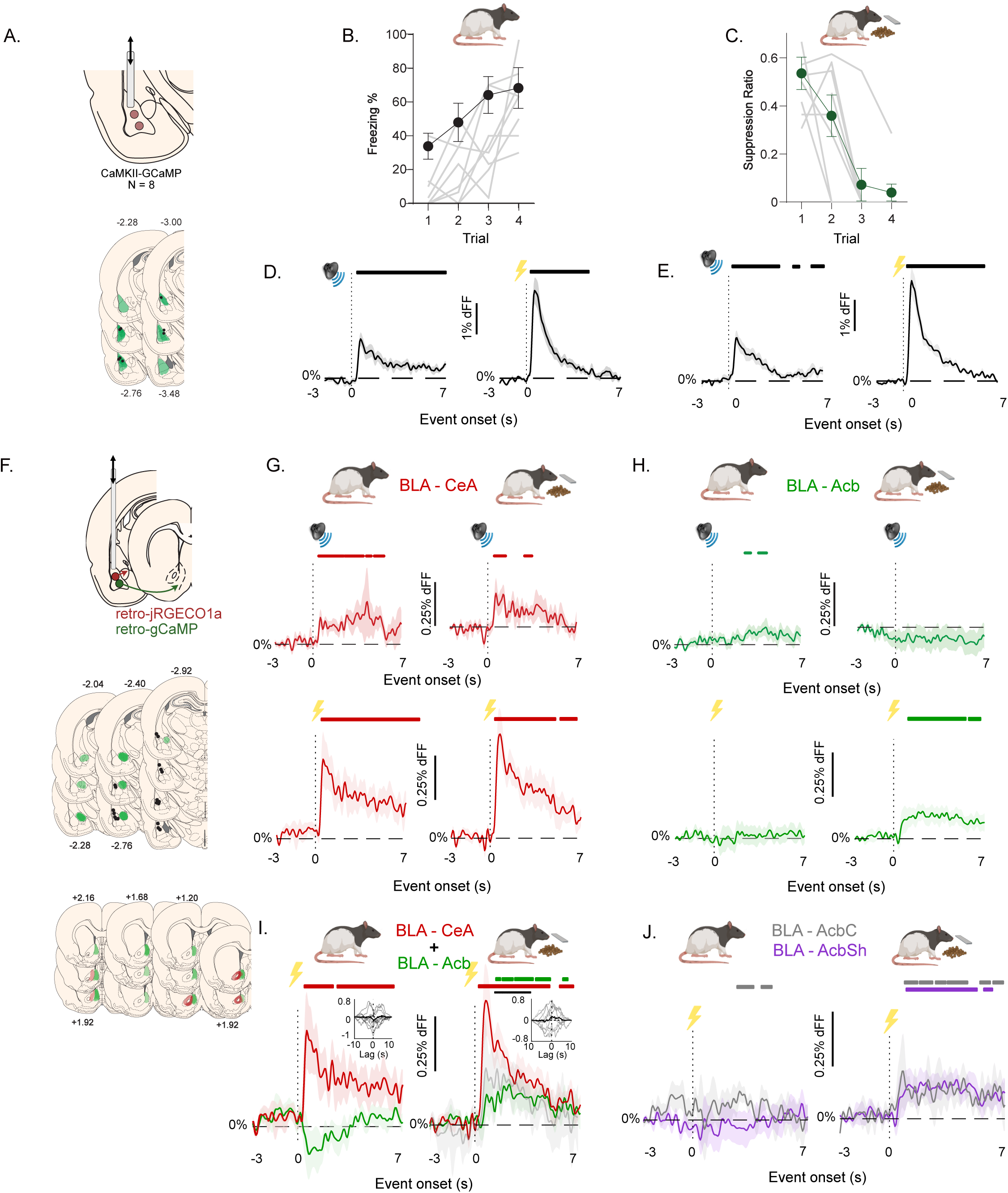
Activity profiles of BLA ➔ CeA and BLA ➔ Acb neurons during fear learning. A) BLA application of CaMKIIa-gCaMP and location of AAV expression (green) as well as fibre tips (black dots) in BLA. B) Acquisition of fear to the auditory CS in the neutral context. C) Acquisition of fear to the auditory CS in the reward context. D). CS- and shock US-evoked transients during fear learning in the neutral context. E). CS- and shock US-evoked transients during fear learning in the reward context. F) Top row. CeA or Acb application of retro-jRGECO1a and retro-gCaMPCaMKIIa-gCaMP. Black dots show location of fibre tips, green shows locations of CeA AAV injections. Middle row. Location of AAV expression in BLA. Bottom row. Location of AAV injections in Acb, green shows injections predominantly in AcbSh, red shows injections predominantly in AcbC. G) CS- and shock US-evoked transients during fear learning in the neutral and reward context for BLA➔CeA neurons. H) CS- and shock US-evoked transients during fear learning in the neutral and reward context for BLA➔Acb neurons. I) Shock US-evoked transients during fear learning in the neutral and reward context for dual colour BLA➔Acb and BLA➔Acb, grey waveform is the within-subjects difference waveform for US-evoked transients. J) Shock US-evoked transients during fear learning in the neutral and reward context for BLA➔AcbC and BLA➔AcbSh neurons. Coloured bars above waveforms indicates periods of statistically significant transients (%DF/F > 0).

Next, we examined the activity of the BLA output pathways across the two different conditions of fear learning. We injected retrograde AAVs containing either gCaMP7f or jRGECO1a into the Acb and CeA and implanted a fibre optic above the BLA to record auditory CS-evoked and footshock US-evoked Ca^2+^-transients in a pathway-specific manner (Figure 3F). BLA➔CeA (n=9) neurons showed robust excitatory Ca^2+^-transients to the auditory CS and footshock US in both the neutral and reward context (Figure 3G). In contrast, BLA➔Acb (n=20) neurons only showed excitatory Ca^2+^-transients to the footshock US in the reward context and showed no detectable Ca^2+^-transients to the footshock US in the neutral context (Figure 3H). BLA➔Acb (n=20) neurons also showed very modest excitatory transients to the auditory CS in the neutral context.

Importantly, we found a similar pattern of footshock US-evoked Ca^2+^-transients when examining the BLA➔CeA and BLA➔Acb pathways within the same subjects (n = 8). Taking advantage of the fact that our within-subject, dual-color photometry permits fair and direct comparison of footshock US Ca^2+^-transients between BLA➔Acb pathway in the same animal in both contexts, we found that footshock US Ca^2+^-transients in the BLA➔Acb pathway were significantly greater in the reward context compared to the neutral context (Figure 3I; grey waveform is difference waveform). Cross-correlation analyses showed that Ca^2+^-transients were modestly anti-correlated in the neutral context but not in the appetitive context (Figure 3I). Finally, we asked whether these profiles of footshock US-evoked Ca^2+^-transients varied across BLA neurons projecting to the core (AcbC) or shell (AcbSh) subregions of the Acb. We divided animals into two groups based on whether AAV injection site predominantly affected AcbC (n=5) or AcbSh (n=13) (Figure 3J). Neither BLA➔AcbC nor BLA➔AcbSh neurons showed robust footshock US-evoked Ca^2+^-transients in the neutral context but both showed robust footshock US-evoked Ca^2+^-transients in the reward context, with the BLA ➔ AcbSh neurons in particular discriminating between the two contexts.

### BLA output pathways have opposing roles in fear learning

Our findings show that BLA neurons respond to a footshock in a circuit- and state-dependent manner. Footshock evoked Ca^2+^-transients in the BLA➔CeA pathway in both the neutral and reward contexts whereas it only evoked transients in the BLA➔Acb pathway in the reward context. Next, we used photoinhibition to determine the roles of this US-evoked activity in fear learning.

First, we first sought to replicate past findings that footshock-evoked activity in a global population of BLA neurons is necessary for fear learning. Rats received CamKIIα-eNpHR3.0 (n = 6) or CamKIIα-eYFP (n = 6), fibre optic implants into the BLA, and we photoinhibited BLA principal neurons at the time of the footshock during fear learning in the reward context (Figure 4A and B). Rats acquired fear to the auditory CS, showing a significant increase in conditioned suppression across days of training (main effect of Day: F(1,10) = 17.57, p = .002). The overall difference in suppression between groups during training approached but did not reach significance (main effect: F (1, 10) = 4.13, p = .069; group x day interaction: F (1, 10) = 4.847, = 0.052). When subsequently tested for fear to the CS without photoinhibition, eNpHR3.0 rats showed significantly less fear compared to controls (main effect of Group: F(1,10) = 11.32, p = .007), including on the first (F(1,10) = 8.345, p = 0.016) and second (F(1,10) = 9.535, p = 0.011) test days. This disruption of fear learning is consistent with previous work in mice (Namburi et al., 2015; Wolff et al., 2014) and rats (Johansen et al., 2014; Sengupta, Winters, Bagley, & McNally, 2016; Sengupta et al., 2018), and shows that the activity of BLA principal neurons at the time of the footshock US supports fear learning.

**Figure 4.**
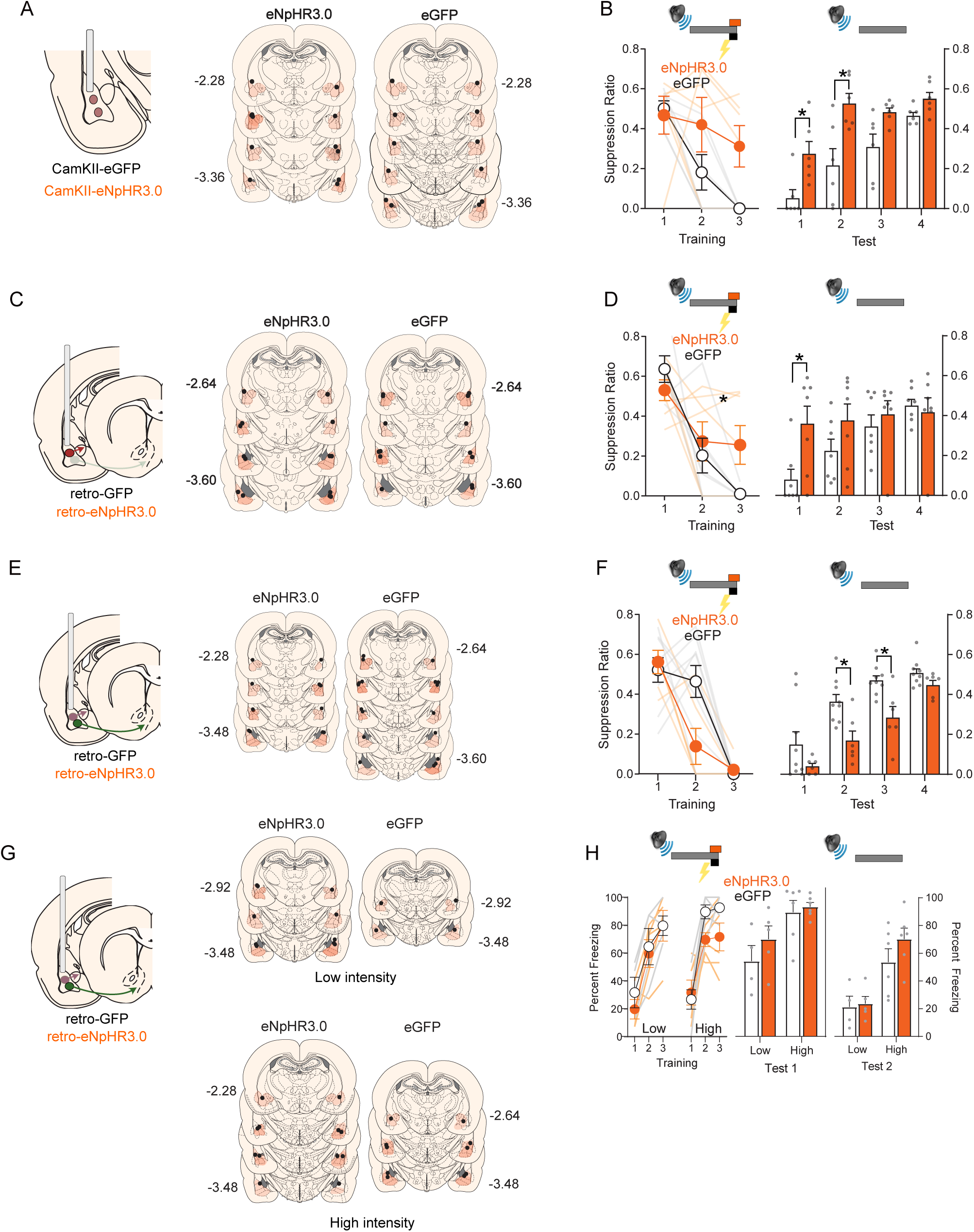
Photoinhibition of basolateral amygdala output pathways and fear learning. A) BLA application of CaMKIIa-eNpHR3.0 or CaMKIIa-eGFP and location of AAV expression (orange) as well as fibre tips (black dots) in BLA. B) Photoinhibition of global BLA neurons during the shock US impaired fear learning in the reward context. C) CeA application of retro-eNpHR3.0 or eGFP and location of AAV expression (orange) as well as fibre tips (black dots) in BLA. D) Photoinhibition of BLA➔CeA neurons during the shock US impaired fear learning in the reward context. E) Acb application of retro-eNpHR3.0 or retro-eGFP and location of AAV expression (orange) as well as fibre tips (black dots) in BLA. F) Photoinhibition of BLA➔Acb neurons during the shock US augmented fear learning in the reward context. G) Acb application of retro-eNpHR3.0 or retro-eGFP and location of AAV expression (orange) as well as fibre tips (black dots) in BLA. F) Photoinhibition of BLA➔Acb neurons during the shock US had no effect fear learning in the neutral context. * p < .05.

Next, we separately tested the roles of footshock-evoked activity in the BLA➔CeA and BLA➔Acb pathways in fear learning in the reward context. We asked how photoinhibition of BLA➔CeA pathway at the time of the footshock affected fear learning. We injected the retrograde AAV containing eNpHR3.0 (n = 7) or the control eGFP (n = 7) into the bilateral CeA and implanted fibre optic cannulae above the BLA (Figure 4C) to silence BLA➔CeA neurons at the time of the footshock US during fear learning (Figure 4D). Rats acquired fear to the CS (main effect of Day: F(1,12) = 33.61, p < .001) and this learning was impaired by BLA➔CeA photoinhibition because the eNpHR3.0 group showed slower rates of fear learning across days compared to controls (Group × Day interaction effect: F(1,12) = 5.13, p = 0.04). Indeed, the eNpHR3.0 group showed significantly less fear to the CS compared to the eGFP group on the last day of fear acquisition (main effect of Group: F(1,12) = 6.33, p = .03). This impairment of learning was also evident when rats were tested for their fear to the CS without photoinhibition. The eNpHR3.0 group showed less fear to the CS on the first day of test compared to controls F (1, 12) = 7.170, p = 0.02). Both groups showed extinction of fear to the CS across the 4 days of test days (main effect of Day: F (1, 12) = 21.06, p = 0.001), but this extinction was faster for the eNpHR3.0 group compared to controls (Group × Day interaction: (F(1,12) = 8.26, p = .01), indicative of less fear learning. So, footshock-evoked activity in the BLA➔CeA pathway drives fear learning.

In a similar manner, we asked how photoinhibition BLA➔Acb pathway at the time of footshock affected fear learning (Figure 4E – F). Rats acquired fear to the CS across learning (main effect of Day: F(1,13) = 121.19, p < .001). The overall difference in suppression between eNpHR3.0 (n = 6) and eGFP (n = 9) groups during training approached but did not reach significance (group main effect: F (1, 13) = 4.54, p = .052; group x day interaction: F (1, 13) = 0.04, p > .05). However, in contrast to the BLA➔CeA pathway, BLA➔Acb photoinhibition augmented fear learning because the eNpHR3.0 group showed significantly more fear to the CS across test compared to controls (main effect group: F (1, 13) = 13.01, p = .003; Day 2 F(1,13) = 10.23, p = 0.007; Day 3 F (1, 13) = 11.29, p = .005). So, footshock-evoked activity in the BLA➔Acb pathway opposes fear learning.

Our dual colour fibre photometry showed that footshock failed to evoke Ca^2+^ transients in the BLA➔Acb pathway during fear learning in the neutral context, in contrast to the robust footshock -evoked Ca^2+^ transients observed in the reward context. This pathway selective activation implies that silencing BLA➔Acb neurons during footshock in the neutral context should have no effect on fear learning. To assess this, we injected the retrograde AAV containing eNpHR3.0 (n = 11) or the control eGFP (n = 10) into the bilateral Acb and implanted fibre optic cannulae above the BLA (Figure 4G) to silence BLA➔Acb neurons at the time of the footshock during fear learning in a neutral context (Figure 4H). We used two different footshock intensities during fear conditioning (0.35mA [n = 9] or 0.7mA [n = 12]) to ensure that any potential effect of fear learning was not masked by a floor or ceiling effect. Rats acquired fear to the CS across learning in both the low footshock intensity (main effect of Day: F(1,7) = 39.99, p < .001) and high footshock intensity (main effect of Day: F(1,7) = 45.36, p < .001) conditions (Figure 4G). However, there was no difference between eNpHR3.0 and eGFP groups in either condition (Low US: main effect of group F (1, 7) = 0.24, p > .05; High US; main effect of group F (1, 10) = 3.28, p > .05). There was also no difference between eNpHR3.0 and eGFP groups across test (Low US: main effect of group F (1, 7) =1.80, p > .05; High US; main effect of group F (1, 10) = 0.93, p > .05). There was significantly more fear on test in the high footshock intensity condition than the low intensity condition (F (1, 17) = 40.81, p < .001), confirming adequate variation in fear levels and statistical power to detect an augmentation of learning by photoinhibition if it had been present.

## Discussion

Here we showed that amygdala circuits are involved in state-dependent opponent processing of fear. First, we showed that BLA➔CeA and BLA➔Acb pathways are largely segregated in the BLA but share overlapping molecular profiles. Using fibre photometry, we showed, for the first time, that BLA ➔ CeA neurons are robustly recruited by footshock regardless of where fear learning occurs whereas the recruitment of BLA➔Acb neurons is state-dependent because footshock only recruited this pathway in a reward context. Finally, using circuit-specific photoinhibition, we showed that this footshock recruitment of BLA➔CeA and BLA➔Acb circuits serves opposing functions in fear learning: activity of BLA➔CeA neurons supports fear learning whereas activity of BLA➔Acb neurons opposes fear learning.

### State-dependent opponent processing of fear

Our key finding is that BLA aversive US processing is circuit- and state-dependent. The BLA➔CeA pathway promotes fear learning. This pathway was recruited by the footshock US regardless of where fear conditioning occurred and silencing this pathway impaired fear learning regardless of where fear conditioning occurred. In contrast, the BLA➔Acb pathway opposes fear learning. This pathway was recruited by the footshock US when fear was conditioned in a reward context, but not neutral context, and silencing this pathway augmented fear learning in the reward but not neutral context. This identification of the BLA➔Acb pathway as a circuit substrate for a footshock US-elicited fear opponent process accords well with findings from rodents and humans implicating the Acb in the relief and safety occasioned by omission or termination of an aversive US (Becerra, Navratilova, Porreca, & Borsook, 2013; Leknes, Lee, Berna, Andersson, & Tracey, 2011; Mohammadi, Bergado-Acosta, & Fendt, 2014). It adds significantly to these findings by showing for the first time that the aversive footshock US can itself activate this fear opponent pathway.

Footshock could recruit the BLA➔Acb pathway in the reward context but not neutral context. The precise driver(s) of this state-dependent recruitment will be important to determine. One possibility is that the reward context increased excitability of BLA➔Acb neurons and hence the likelihood for these neurons to be recruited by the footshock. This is plausible because the BLA➔Acb pathway has a critical role in reward behaviours (Dieterich et al., 2021; Stuber et al., 2011) and increased excitability of BLA neurons increases their likelihood to be recruited during learning (Yiu et al., 2014). This would also be consistent with other evidence that excitability of BLA➔CeA and BLA➔Acb neurons is modulated by recent experiences, including learning and changes in internal states. For example, fear conditioning strengthens the BLA➔CeA pathway but weakens the BLA➔Acb pathway whereas reward learning has the opposite effect (Namburi et al., 2015). Food deprivation increased baseline activity of BLA➔Acb neurons but decreased baseline activity of BLA➔CeA neurons. Furthermore, food deprivation switched the relationship between BLA➔CeA and BLA➔Acb neurons, with BLA➔ CeA photostimulation inhibiting BLA➔Acb neurons in non-deprived mice but exciting BLA➔Acb neurons in food deprived mice (Calhoon et al., 2018). These findings show that interactions between the BLA➔CeA and BLA➔Acb pathways are not invariant and are consistent with our finding of circuit- and state-dependent opponent processing of fear.

It is worth emphasizing that food restriction *per se* is not the critical variable dictating recruitment of the BLA➔Acb pathway by footshock. We used a within-subjects design to control this possibility, with concurrent measurement of the BLA➔CeA and BLA➔Acb pathways in the same animals under the same food restriction regimes. Regardless, understanding precisely how and when the fear opponent role of the BLA➔Acb pathway is unmasked could prove useful in designing novel interventions to minimize the amount of fear learned after a traumatic experience.

Interestingly, none of the molecular markers we used (*Rspo2*, *Ppp1r1b* and *Fezf2*) were able to distinguish fear opponent BLA➔Acb neurons from fear promoting BLA➔CeA neurons. This was surprising because these markers have proved useful in delineating valence coding neurons in the mouse BLA but were co-expressed in rat BLA➔Acb and BLA➔CeA neurons. It is possible that the US-elicited fear opponent process identified here is distinct from, or due to a subset of, valence coding BLA neurons. One marker that could be worth examination is *Thy-1*. Excitation of *Thy-1* expressing BLA neurons at the time of the footshock US inhibits fear learning (Jasnow et al., 2013) – a functional profile consistent with a fear opponent process - and BLA *Thy-1* neurons also project to Acb (Porrero, Rubio-Garrido, Avendaño, & Clascá, 2010). So, BLA *Thy-1* neurons may be a cellular-substrate of the fear opponent-process discovered here.

### Methodological considerations

There are two issues to consider. First, we used dual colour photometry via jRGECO and GCaMP to assess footshock US-evoked activity in BLA➔CeA and BLA➔Acb neurons. A key consideration is spectral separation of these signals. It is unlikely that a lack of spectral separation can explain our findings: we counterbalanced which calcium indicator was expressed in the two pathways, we recorded during the same footshock events in the same animals in the reward and neutral context, and the state- and circuit-dependent effects of photoinhibition supported findings of state and circuit-dependent recruitment. Second, our experiments were restricted to male animals due to unavailability of female animals. It is important to note that many of our findings (global BLA footshock-evoked transients, effects of global BLA silencing, effects of BLA➔ CeA silencing) recapitulated past work in both sexes in rats and mice, so there are strong grounds for assuming generality of our findings. However, given that there are sex differences in activity of amygdala neurons linked to differences in excitatory synaptic input (Blume et al., 2017) this will remain an important empirical question.

### Conclusions

Here we show that BLA aversive US processing is circuit- and state-dependent. Whereas the BLA ➔ CeA pathway was recruited by the aversive footshock US regardless of where fear learning occurred, the BLA➔Acb pathway was recruited by the footshock US only in the reward context. This footshock-driven activity in the BLA ➔ Acb pathway functionally opposed the BLA ➔ CeA pathway to limit how much fear was learned.

## Acknowledgements

Author Contributions: <u>Conceptualisation</u>: GPM, JOYY, AL; <u>Methodology</u>: JOYY, AL, LL, GPM; <u>Investigation</u>: AL, JOYY, LA; <u>Writing – Original Draft</u>: JOYY, GPM. <u>Writing – Review & Editing</u>: All authors. <u>Funding Acquisition</u>: GPM. This work was supported by an Australian Postgraduate Award, the National Health and Medical Research Council (GNT2011633), and the Australian Research Council (DP220100040).

